# How to catch flies in the city (fast): Citizen Science on *Drosophila* ecology helps to raise awareness for biodiversity in urban environments

**DOI:** 10.1101/2025.10.10.681315

**Authors:** Isolde Gottwald, Sonja Steindl, Flora Strasser, Megumi Kiesel, Heimo Rainer, Elisabeth Haring, Martin Kapun

**Author notes:** co-correspondence. co-shared first authors (Isolde Gottwald, Sonja Steindl).

## Abstract

To efficiently mitigate biodiversity loss both robust ecological data and broader societal engagement are needed. Citizen science offers a pathway to address these dual challenges by combining data collection with public involvement. Here, we introduce the citizen science project *Vienna City Fly,* conducted at the Natural History Museum Vienna (NHMW), which used fruit flies (*Drosophila*) as a model organism group for urban biodiversity research and public engagement. Participants deployed standardized traps across Vienna, enabling systematic sampling and active participation in research, which resulted in hundreds of fly collections, thousands of sampled fly specimens and new insights into the biodiversity and ecology of the collected fly species, demonstrating the feasibility of large-scale ecological research through public collaboration. To assess motivational drivers and perceived impacts of participation, we conducted an online survey (N=59) using the validated psychometric MORFEN-CS scale. Survey results revealed that nature conservation values and the intention to contribute to biodiversity conservation emerged as the strongest drivers of engagement, complemented by sociopolitical responsibility and citizen science-based motivations. Reported outcomes include knowledge gain (61%), more positive attitude toward study organisms (50%), and increased awareness of biodiversity (30%). Satisfaction was very high, with 85% rating their experience at the top of the scale and all participants expressing willingness to join future citizen science projects. Recruitment occurred mainly via social networks, and the sample of participants was highly educated, indicating limits to inclusivity and reach. Our findings demonstrate that citizen science can contribute to robust ecological data collection while gaining knowledge and awareness. Natural history museums, as trusted institutions, play a key role in facilitating such initiatives. Overall, *Drosophila* research proves to be a suitable field for citizen science, combining accessibility with strong potential for advancing ecological and biodiversity research as well as public engagement in urban biodiversity research.

## INTRODUCTION

Biodiversity loss is widely recognized as one of the most pressing environmental and societal challenges of our times (IPBES 2019). It directly impacts ecosystem stability, as well as human health, food security and well-being (Cardinale et al. 2012), and is therefore a central challenge that the United Nations Sustainable Development Goals (SDGs) seek to address. In light of the accelerating ecosystem degradation driven by human activity, there is an urgent need to collect data on the ecosystem stability and biodiversity loss (Hawkins 2024) and at the same time foster collective societal responsibility and enable the behavioral transformations necessary for biodiversity protection (Nielsen et al. 2021). On top of that and in light of a growing disconnect from the natural world (Beery et al. 2023), fostering public engagement in biodiversity research and awareness of nature and its relevance should be of high priority (Greving et al. 2022).

In this context, citizen science – defined as the active participation of non-professionals in scientific research – has emerged as a promising approach to generate valuable data and enhance societal engagement with science (Beck et al. 2020). Citizen science is also gaining recognition in the context of the Sustainable Development Goals (SDGs), particularly biodiversity (SDG 15 – life on land), although its role is only beginning to be systematically acknowledged (Fraisl et al. 2020, Moczek et al. 2021a). For example, citizen science may combine environmental data collection with public participation (van Noordwijk et al. 2021) and advance knowledge production. It may foster environmental awareness, build scientific literacy (Liu and Kobernus 2017, Sauermann et al. 2020), and may enable civic engagement, supporting evidence-based decision-making (Sauermann et al. 2020, Turrini et al. 2018) and appreciation of nature and social engagement (Hecker et al. 2018) – all critical in light of the globally accelerating biodiversity loss (Hawkins 2024). Beyond advancing research, citizen science strengthens ties between science and society (Serrano Sanz et al. 2014). Citizen science also promotes active citizenship, motivates socio-political change (Hecker et al. 2018), and encourages behavioural shifts towards sustainability (Sauermann et al. 2020). Citizens thus play crucial roles as consumers, advocates and voters, positioning their engagement as vital for sustainability transitions (Sauermann et al. 2020). Citizen science further advances conservation – from ecological monitoring and species tracking to other forms of biodiversity documentation (Ballard et al. 2017, Greving et al. 2022). In summary, citizen science is well established in the natural sciences (Frigerio et al. 2021), particularly in environmental and biodiversity research (van Noordwijk et al. 2021).

Motivations for participation include gaining knowledge and awareness, contributing to the common good, protecting the environment, and supporting conservation initiatives – often reinforced by recognition of the contributions from participants (Asingizwe et al. 2020, Hobbs and White 2012, Kühn et al. 2022). Additional drivers include personal empowerment, a deeper appreciation of nature, increased social engagement, and willingness to act as environmental opinion leaders (Hecker et al. 2018, Liu and Kobernus 2017). Recent studies report increased knowledge, more positive attitudes towards the objects studied and enhanced scientific skills among participants (e.g. Aristeidou and Herodotou 2020, Greving et al. 2022, Kühn et al. 2022, Lynch et al. 2018, Moczek et al. 2021b, Toomey and Domroese 2013).

Natural History Museums (NHMs) illustrate the science-society interface particularly well. Combining the preservation of collections with the exchange of knowledge, long tradition of working with amateur naturalists, expertise in science communication, and a trusted public profile, NHMs are uniquely positioned to facilitate citizen science (Ballard et al. 2017, Lindner et al. 2024, Sforzi et al. 2018). NHMs help to involve the public in research, increase awareness of environmental change, and support conservation efforts (Sforzi et al. 2018). Often located in city centers, they have a unique power to engage residents and visitors in regional and urban biodiversity topics (Ballard et al. 2017). Examples range from gains in scientific knowledge and an increased place connectedness (Ballard et al. 2017), combining citizen data with museum collections to track species responses to climate change in Denmark (Olsen et al. 2020) to large-scale documentation of invasive species in Japan (Ishida 2020). In Europe, several platforms such as in Germany (https://www.mitforschen.org/), and in Austria (https://www.citizen-science.at/) illustrate the institutionalization of citizen science (Haklay et al. 2021). Still, recent findings from the Austrian Science Barometer (Austrian Academy of Sciences 2024) highlight a growing disconnect between science and society: Public interest in science has declined to 56%, and only 32% of respondents feel well informed. This underscores the need for new, more inclusive forms of public engagement that extend beyond conventional science communication. To explore this potential, we employ *Drosophila* science as a novel and powerful model system for participatory research. Importantly, our approach extends beyond the classical laboratory model species to include all *Drosophila* species occurring within the study area, thereby linking urban ecological research with broader biodiversity perspectives. This dual focus not only generates valuable data but also provides an innovative platform for raising awareness and engaging the public in urban biodiversity research.

Species of the genus *Drosophila* – commonly referred to as “vinegar flies” or “fruit flies” – are small dipterans that occupy a wide variety of ecological niches. Several generalist species are closely associated with humans and human activity, particularly those belonging to a group of synanthropic taxa collectively referred to as the *worldwide guild* (Atkinson and Shorrocks 1977, Miller et al. 2017, Nunney 1996). These species with broad ecological tolerance are globally distributed, frequent visitors to human dwellings, and are often regarded as pests or, at the very least, nuisance organisms by the general public (Markow 2015). In particular, *D. melanogaster*, one of the best-studied genetic model organisms, belongs to this group of generalist species (Bilder and Irvine 2017, Haudry et al. 2020, Keller 2007, Lachaise et al. 1998). In contrast, other *Drosophila* species are strict ecological specialists with narrow dietary and habitat preferences and correspondingly restricted distributions, often limited to environments with little human influence. With approximately 1,600 species described worldwide (Bächli 1982, Brake and Bächli 2008, O’Grady and DeSalle 2018), the genus *Drosophila* thus represents a versatile and promising system of bioindicators, potentially allowing assessments of the degree of human disturbance in ecosystems based on species composition (Parsons 1991, Poppe et al. 2013).

Moreover, many *Drosophila* species are easy to sample and, due to the strong association of some species with humans, they are well-suited for citizen science projects, which not only engage public helpers in data collection but also provide education about biodiversity, biodiversity loss, and the biology and natural history of the studied species. Several successful initiatives have already demonstrated the potential of this approach. For example, *Melanogaster: Catch the Fly* (https://melanogaster.eu/) an originally Spanish project, effectively involves school pupils in sampling –preferentially *D. melanogaster –*while teaching the students about evolution and insect diversity in rural environments.

*Drosophila* fruit flies thus provide a suitable model for assessing large-scale human-driven processes such as urbanization and their impact on biodiversity in the face of ongoing environmental changes such as the global climate change (Parsons 1991, Rodríguez-Trelles et al. 1998, Rodríguez-Trelles and Tarrío 2024). As a major driver of environmental change, urbanization alters biodiversity through processes such as habitat fragmentation (Gibb and Hochuli 2002), habitat loss (Czech et al. 2000), urban heat island effects (Schwaab 2022), noise pollution (Sordello et al. 2020), and increased human proximity. Quantifying the responses of *Drosophila* communities to these pressures requires dense spatial sampling, which can be efficiently implemented through citizen science initiatives. Besides generating comprehensive data for fundamental research, citizen science projects also help bridging the critical gap between science and public understanding of urban biodiversity (Theobald et al. 2015). Fruit flies are abundant in households and therefore readily tractable, yet their species richness and ecological diversity in urban areas is largely unknown to research and the general public. Instead, they are often perceived negatively or regarded as biologically uninteresting. This combination makes them a particularly suitable model system for investigating biodiversity in ephemeral urban environments, while also providing an opportunity to examine how participation in fruit fly–based citizen science projects influences public perception of insects and biodiversity awareness.

To address these largely unexplored opportunities, we initiated the citizen science project *Vienna City Fly* (VCF) in the metropolitan area of Vienna. The overarching goal of this project was to establish a structured citizen science research program and conceptual framework that uses *Drosophila* both as a model for investigating biodiversity and ecological patterns in urban environments and as an educational tool to raise biodiversity awareness.The biological results and ecological analyses of the survey on *Drosophila* species in the urban environment of Vienna are described in details elsewhere (Kapun et al. 2025).

Understanding why individuals participate in citizen science is key for attracting and retaining participants, ensuring data quality, and enhancing institutional acceptance (Geoghegan et al. 2016, Land-Zastra et al. 2021). Despite its centrality to project effectiveness, research on motivational drivers and the impact of participation on attitudes, behaviour and knowledge remains limited (Kam et al. 2021; Somerwill and Wehn 2022, West and Pateman 2016), partly due to inconsistent methodologies and a lack of standardized instruments (Greving et al. 2022, Kühn et al. 2022, Moczek et al. 2021b). Incorporating methods of social sciences and environmental psychology can address these gaps and enrich understanding of citizen science outcomes (Tauginienè et al. 2020).

To this end, this study reports the results of an online survey among the participants of the Citizen Science project *Vienna City Fly* centered around urban *Drosophila* and employs the validated psychometric *Motivational and ORganisational Functions of voluntary ENgagement in Citizen Science* scale (MORFEN-CS), measuring motivational and organizational functions of volunteering in biodiversity and environmental sciences. This instrument builds on the theoretical Model of Influences on Participation in Citizen Science (Geoghegan et al. 2016, West and Pateman, 2016) and Penner’s Model Causes of Sustained Volunteerism (Penner 2002) as well as prior related inventories (see Figure 2, Moczek et al. 2021b). By integrating insights from psychology and sustained social volunteerism, MORFEN-CS provides a theoretically grounded and psychometrically validated instrument to systematically assess motivational and organizational functions driving participation in citizen science projects. Previous applications of the scale, such as in the *Insects of Saxony* project, highlight the salience of prosocial motivations – particularly contributing to nature conservation in socially and politically meaningful ways – alongside factors such as communication, feedback and organization, which sustains participation (Moczek et al. 2021b). Participation is also motivated by opportunities to understand scientific processes and engage in communities with shared (research) interests, with – consistent with previous studies – altruistic motivation rated higher than self-oriented motives (Moczek et al. 2021b). The scale was adapted accordingly (see Methods and Appendix 3).

**Figure 1.**
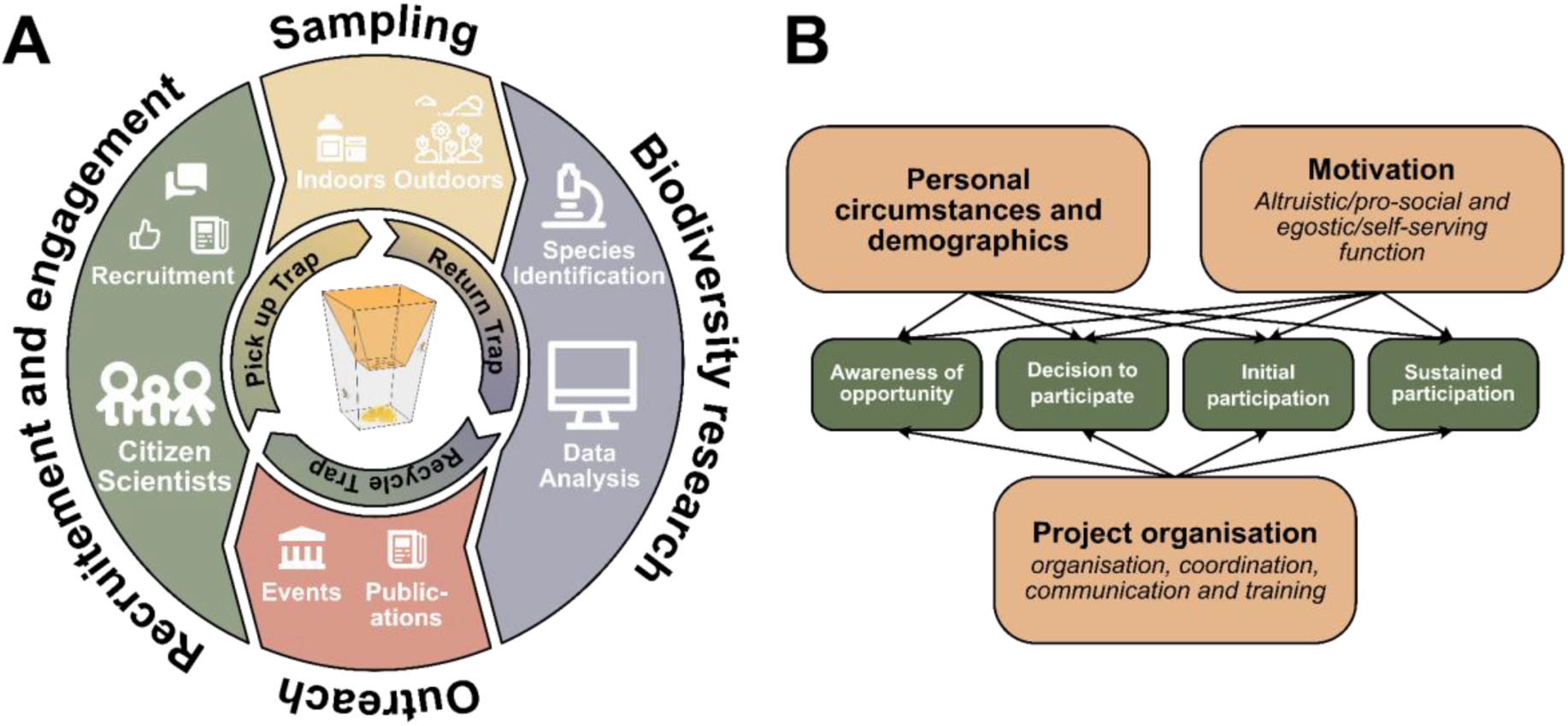
(A) Concept of the citizen science project *Vienna City Fly*, which is structured around four main pillars: (1) recruitment and engagement of citizen scientists through diverse communication channels, (2) sampling of *Drosophila* specimens indoors and outdoors by citizen scientists, (3) biodiversity research through data analysis of collected specimens, including species identification and ecological inference based on species abundances across locations, and (4) dissemination and outreach through museum events, media publications, and interactions with citizen scientists. The inner circle, enclosing a schematic funnel trap with bait and flies used in this project, illustrates the rotation scheme in which citizen scientists collect empty traps at the Natural History Museum Vienna, deploy them in their homes, and return filled traps in exchange for new ones. (B) Model of Influence for Participation in citizen science projects by Penner (2002), adapted by and cited from West et al. (2016) in Geoghegan et al. (2016). Additions by Moczek (2019), as cited in Moczek et al. (2021b)

**Figure 2.**
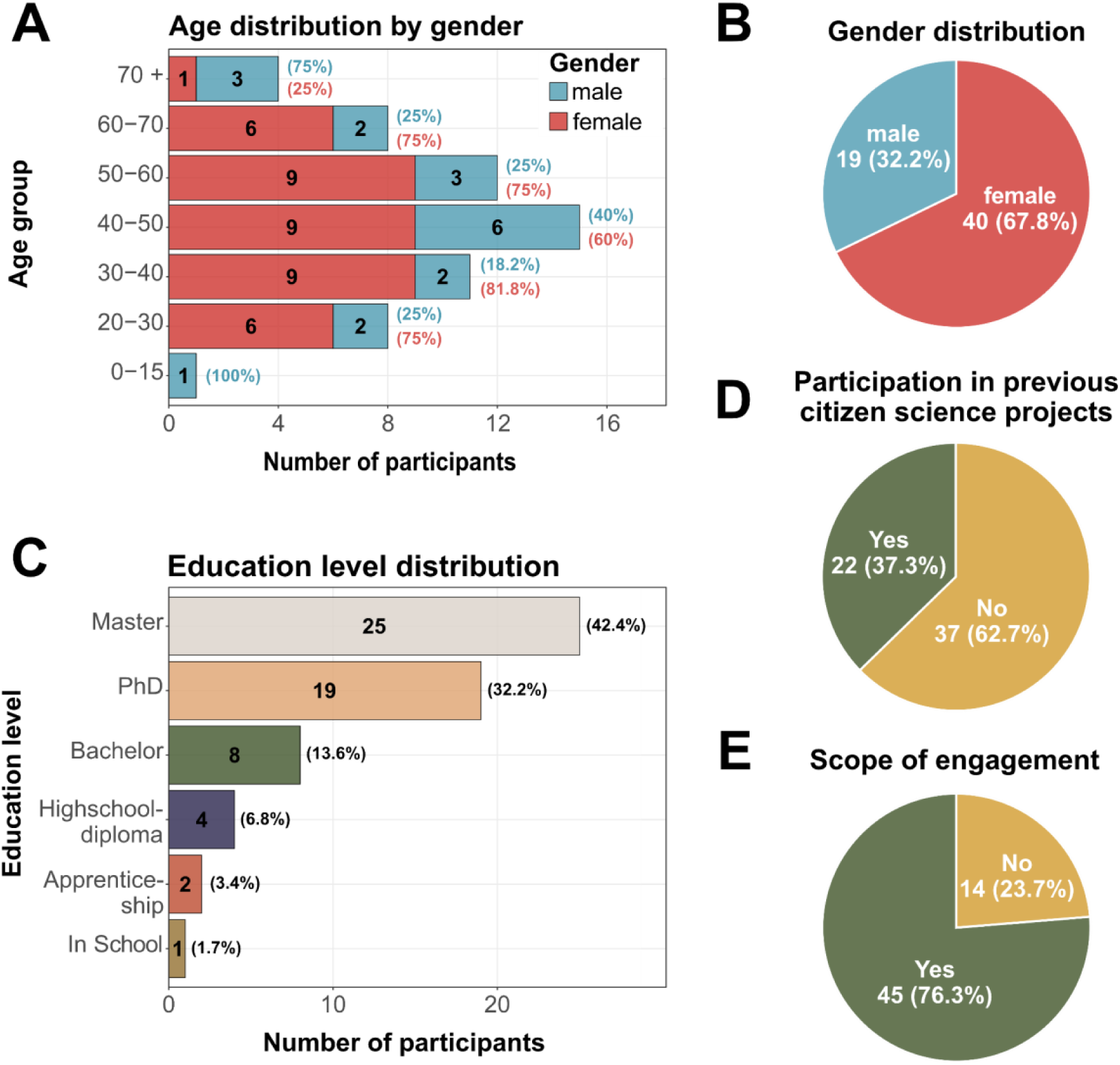
Bar– and piecharts summarizing sociodemographic variables of the citizen science participants: (A) Age distribution by gender, (B) Gender distribution, (C) Education level distribution, (D) Participation in previous citizen science projects, and (E) Scope of engagement, i.e., if flies were successfully sampled.

In summary, in this study, we present methodology developed to engage and interact with citizen scientists using *Drosophila* research to evaluate how participation in the project influences their awareness of, and interest in, biodiversity and scientific research.

## METHODS

### Recruitment strategies and *Drosophila* sample collection

The citizen science project *Vienna City Fly* was initiated as a grass-roots effort to collect species of the genus *Drosophila* in urban areas, particularly within homes and gardens, with the help of citizen scientists and which employed multiple recruitment strategies to engage participants (see Figure 1). Recruitment was conducted through (1) personal and professional communication within the social network of the Natural History Museum Vienna (NHMW), (2) social media campaigns on the official Facebook, Instagram, and YouTube channels of the NHMW (including the production of an explanatory video), (3) newspaper articles in the Austrian press, and (4) a dedicated project webpage providing information and participation guidelines (https://nhmvienna.github.io/ViennaCityFly/). In addition, (5) participants were directly recruited at NHMW outreach events, where the project was presented by the initiators at science booths.

As our research focused on assessing the biodiversity and ecology of *Drosophila* communities in urban and suburban environments, we concentrated recruitment efforts on citizen scientists residing within or near Vienna with approximately 2 million inhabitants. Vienna is the capital and largest city of Austria, covers more than 414 km² and exhibits typical Central European urban characteristics (Bauer et al. 2024). These include a densely built historic center surrounded by less dense residential areas, extensive infrastructure, and heterogeneous land-use types ranging from residential and industrial zones to forests and agricultural land. Vienna is traversed by approximately 20 km of the Danube River. Administratively, the city is divided into 23 districts. To capture variation in urbanization intensity, we recruited citizen scientists from as many districts as possible, thereby representing a broad gradient of urban conditions. In addition, participants from surrounding rural villages in neighboring federal states, dominated by agricultural land use, were engaged to complement the urban dataset.

To facilitate participation, we established an online registration webpage where prospective citizen scientists could provide information such as the intended sampling location, sampling period, and contact details. In compliance with the General Data Protection Regulation (GDPR) of the European Union, participants were assigned pseudonymized identification numbers, which were subsequently linked to the corresponding sampling coordinates. To ensure standardized and straightforward fly collections, participants were provided with commercial funnel traps. These traps consisted of a transparent outer container holding an opaque conical funnel with three small holes at the bottom (see center of Figure 1). Slices of fruit, such as lemon, banana, or apple, were placed at the base of the trap to serve as bait. Flies entered through the holes at the bottom of the funnel and were retained inside due to behavioral constraints, including negative geotaxis (upward movement) and positive phototaxis (movement toward light), which prevented specimens from locating the exit. To facilitate successful *Drosophila* collections, participants were provided with detailed sampling instructions and were asked to place the traps for a maximum of 14 days either indoors (e.g., in the kitchen or living room) or outdoors (e.g., on a balcony or in a garden) in locations protected from rain. Trap distribution and collection were organized through a rotation system coordinated with the service staff of the NHMW. Citizen scientists picked up empty traps at the museum and returned filled traps along with metadata, including the type of bait used, the trap location (indoors or outdoors), and the duration of trapping, in exchange for new empty traps (see inner circle of Figure 1A). To further encourage participation, each volunteer received a small sample of honey produced on the roof of the NHMW as a token of appreciation upon returning a filled trap. These measures ensured standardized sample collection and provided a basis for engaging participants throughout the project.

### Dissemination strategies and assessment of the participant motivation

We employed several strategies to interact and communicate with citizen scientists. A dedicated email address was established to allow direct communication and to answer questions regarding the sampling methods. Regular digital newsletters were sent to keep participants informed about the progress of the project, and outreach activities at the NHMW provided opportunities for personal contact and for reporting preliminary results. After the project concluded in December 2024, all participants were invited to a workshop at Deck 50 of the NHMW, an open and participatory laboratory space. During the workshop, which took place in January 2025, we presented scientific analyses and results derived from the data collected, and offered a hands-on session in which participants could observe fruit flies under a microscope and practice distinguishing different *Drosophila* species independently. The scientific results of the project were further disseminated through newspaper articles, a television documentary broadcast on the Austrian public TV station ORF1 and a scientific publication (Kapun et al. 2025).

### Data collection and measures

Finally, we conducted an online survey in German among all participants to assess the primary motivations for taking part in our citizen science project and to investigate whether participation influenced the perception of science in general – particularly awareness of biodiversity, loss of biodiversity, and the biology of fruit flies. In this survey, participants provided age, gender and highest level of education after giving informed consent. To measure engagement, we assessed duration and frequency in previous citizen science projects as well as scope of engagement in the current project, i.e., whether *Drosophila* specimens were successfully sampled. After this, the MORFEN-CS scale was employed, followed by a self-assessment of the impact, overall assessment of their contentment with the project as well as intended future involvement. The MORFEN-CS scale is divided into multiple subscales, each assessing different aspects of this type of motivation. Six of these subscales were selected and adapted for the purpose of this project as considered most relevant to its objectives and aligned with the most strongly expressed subscales identified by Moczek et al. (2021b) (see Supplementary Materials). These include Nature Conservation Values, Social Motives, Sociopolitical Responsibility, Citizen Science, Communication and Organization. From the latter two subscales, one Communication item and three organization items were selected and combined into a single Communication-Organization subscale. Furthermore, an additional question was formulated to assess curiosity about the subject of the study (“I am involved in the project because I want to learn more about the distribution of fruit flies and their way of life.”). All items were measured on a 5-point Likert scale ranging from 1 (completely disagree) to 5 (agree completely). To tailor the questionnaire to the needs of our target group, we adapted minor formulations of the original scale (for the full survey and documented adaptations to the MORFEN-CS scale, see Supplementary Materials and Table S1). The survey ended with an option to provide feedback and a debriefing about data protection.

### Data analysis

Statistical analyses were performed in R (v. 4.2.1; R Core Team 2019) with the help of Microsoft Co-Pilot (LLM Model: Claude Sonnet 4 from Anthropic) for code documentation, auditing and refactoring following manual data cleaning in Microsoft Excel. Demographic variables were summarized and visualized using functions from the tidyverse package in R, specifically dplyr() for data manipulation and ggplot2() for data visualization (Wickham et al. 2019). Six motivation indices were calculated from Likert-scale responses: Nature Conservation Values (NCV_idx; 5 items), Social Motives (SM_idx; 3 items), Sociopolitical Responsibility (SPR_idx; 3 items), Citizen Science Interest (CS_idx; 5 items), Communication & Organization (CommOrg_idx; 4 items), and Curiosity to Learn More (SPR_extra; 1 item). Each composite index was computed as the mean of constituent items, requiring a minimum of 50% item availability. An overall motivation index (MeanMotivationOverall) was derived from all subscales except SPR_extra. The internal consistency of each multi-item subscale was assessed using Cronbach’s alpha with the psych package (Revelle, 2022), following established reliability thresholds (α ≥ 0.9 = excellent; 0.8-0.89 = good; 0.7-0.79 = acceptable; 0.6-0.69 = questionable; <0.6 = poor). Descriptive statistics for motivation scales included means, standard deviations, and distributional characteristics visualized through boxplots with ggplot2(). Inter-scale relationships were examined using Pearson correlations with significance testing and visualized using correlation matrices from the corrplot package (Wei et al. 2021). We further performed one-way ANOVA to test for differences between motivation scales, with Bonferroni-corrected post-hoc pairwise comparisons conducted using the rstatix package (Kassambara 2021) when omnibus tests were significant (p < 0.05). Eta-squared (η²) was calculated as a measure of effect size using base R statistical functions. Following the one-way ANOVA, we computed eta-squared as the ratio of between-group sum of squares to total sum of squares (SS_between / SS_total) to quantify the proportion of variance in motivation scores explained by differences between motivation scales. Values of 0.01, 0.06, and 0.14 represent small, medium, and large effect sizes, respectively (Cohen 1988). Finally, we analyzed engagement outcomes (project impact, satisfaction, and future participation intentions) by parsing comma-separated responses, using the dplyr() function to accommodate multiple-choice answers. Frequencies and percentages were calculated with dplyr() and the results were visualized with ggplot2(), respectively.

## RESULTS

In this project, we developed a citizen science programme to study the distribution and ecology of urban *Drosophila* while simultaneously fostering biodiversity awareness and public interest in science by involving citizen scientists in fruit fly collection. Our recruitment and communication strategy as well as the rotation scheme for picking up and returning fly traps as described in the “Methods” section was well received by a total of 163 recruited participants, out of which 89 actively provided flies for the project.

With our method, which can be easily adopted for similar future citizen scientist projects, we were able to obtain a total of 18,173 flies in 279 traps that were collected within a six month sampling period ranging from June 2024 to December 2024. The collection comprised thirteen *Drosophila* species, including *D. mercatorum* and *D. virilis*, which were documented for the first time in Austria (Kapun et al. 2025). By combining abundance data from multiple *Drosophila* species with comprehensive climatic information, we were able to generate robust predictions about ecological differences among *Drosophila* species in urban environments across the Vienna metropolitan area (Kapun et al. 2025). Overall, these findings underscore the value of citizen science for biodiversity research – particularly because dense sampling over such a large area, which is essential for reliable ecological inference, would not have been possible without the contribution of numerous volunteers.

### Demographics of citizen scientists

To better understand who participated in the project and what motivated their involvement, we conducted a demographic and motivation survey among the registered citizen scientists between December 21^st^ 2024 and January 14^th^ 2025. Out of the 163 registered citizen scientists, a total of 63 (38.6%) participants consented to take part in the survey; however, only 59 (36.2%) were included in the final analysis, as four participants reported being unable to fully contribute to the project (e.g., due to not having caught any flies). Approximately 40% of the participants were older than 50 years, a quarter (25.4%) between 40 and 50 years and 33.9% younger than 40 (Figure 2A). Two thirds of the citizen scientists were female (66.7%; Figure 2B) and a majority of the respondents had a master’s degree (42.4%) or higher (32.2%; see Figure 2C). 37.3 percent had previously participated in citizen science projects (Figure 2D). 76.3% fully contributed to the project until its conclusion by providing flytraps (Figure 2E).

We further evaluated the efficiency of our recruitment methods and found that most participants (71%) learned about the project through their social networks, while 14% were informed via media channels and 6% through museum events (Figure 3A).

**Figure 3.**
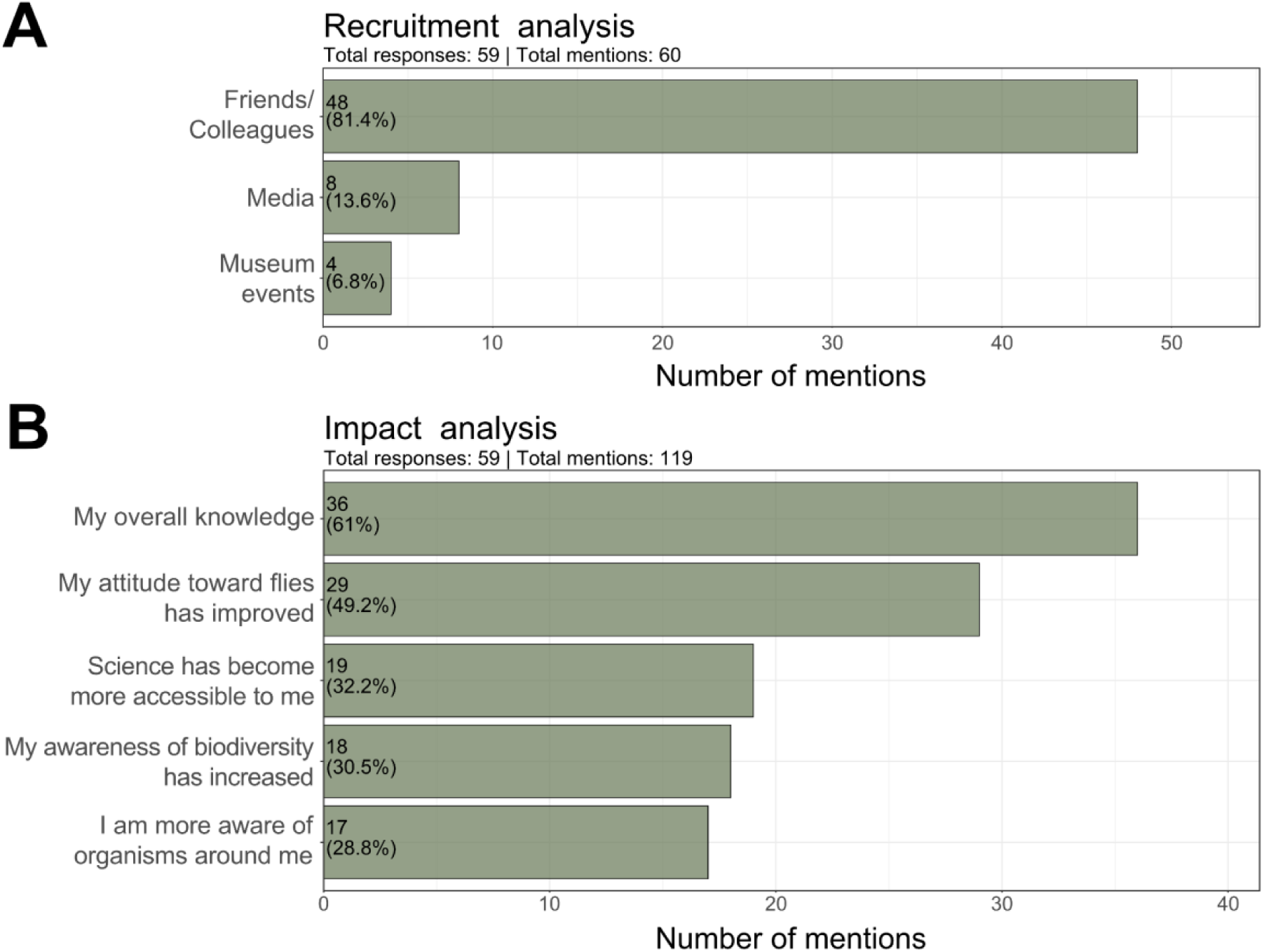
Barplots showing the proportion of responses to the questions “How did you hear of the project?”(A: Recruitment of participants) and “What has changed as a result of participating in the project?”(B: Self-reported impact of participation)

### Perceptions of engagement impact

To assess level and type of engagement, participants were asked whether their involvement had any impact – essentially a form of self-reported impact assessment – with six response options available (multiple answers could be selected, results see Figure 3B). The most frequently chosen response was that their knowledge had improved: 61% of respondents stated that their participation had increased their knowledge. Additionally, 49% reported a positive shift in their attitude toward the study subject. The second most common response, selected by 32% of participants, was that science had become more tangible through their involvement. Furthermore, 31% indicated an increased overall awareness of biodiversity, while 29% noted a heightened awareness of organisms in their immediate environment. Notably, no participant reported a worsening of their attitude toward the study subject.

When asked for suggestions for improvement, 43% indicated interest in receiving more scientific content for discussion. Additionally, 10% of respondents suggested enhancements in project logistics, and 8% desired improved communication from the project organisers. Many participants used the open-ended response option to express overall satisfaction and offered no suggestions for improvement. The final feedback was predominantly positive and included praise for the project implementation, interest in participating in similar future projects, and enthusiasm for the results. Some feedback also included constructive suggestions for further improvement. A large majority of respondents (85%) expressed high satisfaction with their participation, giving a rating of 4 or 5 on a 5-point Likert scale (Figure 4A). Only one person reported dissatisfaction. All respondents expressed a desire to continue participating in citizen science projects (100%).

**Figure 4.**
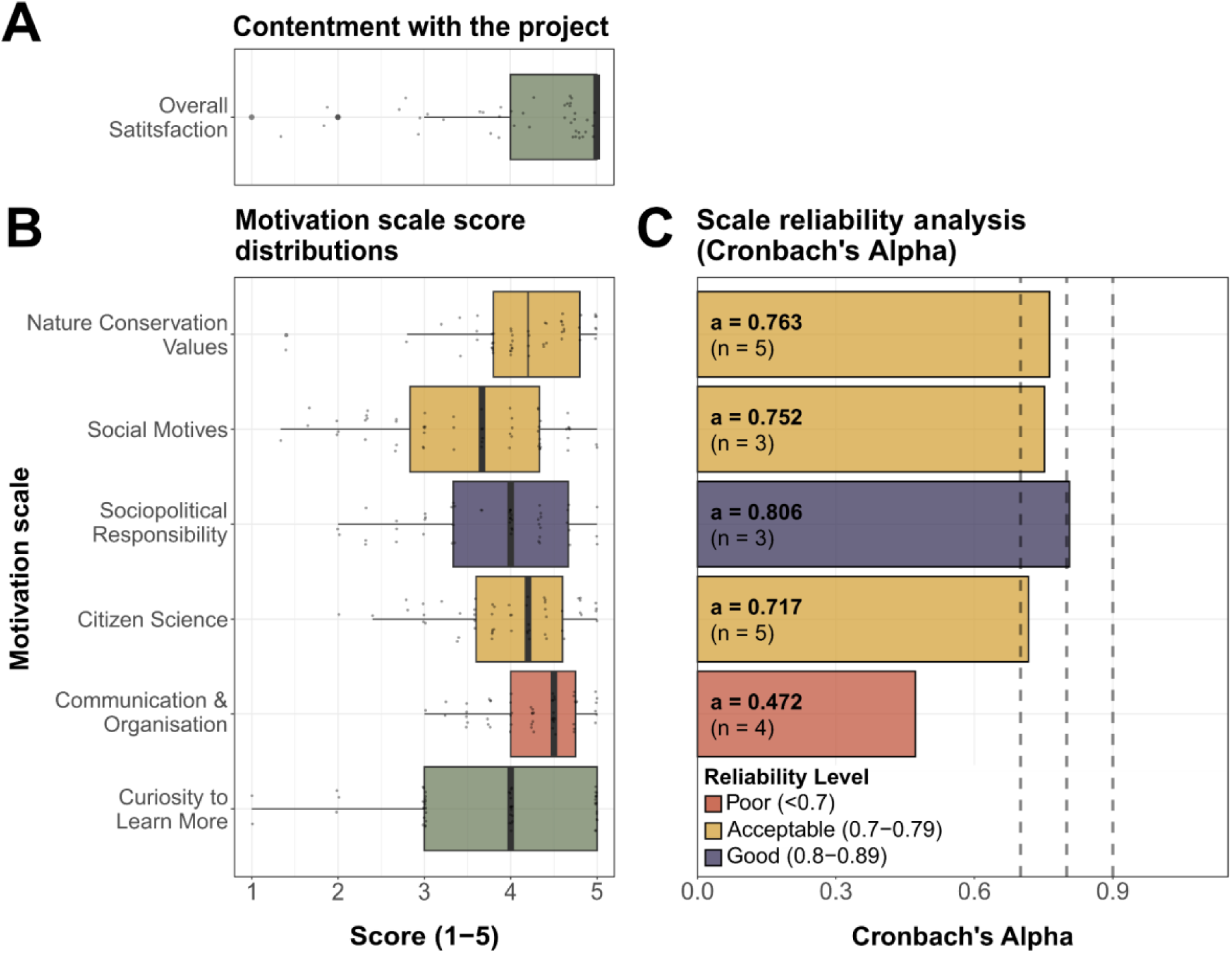
Boxplots show Linkert-scale scores for (A) contentment with the project and for (B) different MORFEN-CS motivation subscales. Panel C depicts the corresponding reliability analyses for five of the six subscales. The boxplots show the median as well as interquantile ranges.

### Motivation

The strongest motivational factor was Communication and Organization (M = 4.29 (SD = 0.53), closely followed by Nature Conservation Values (M = 4.24, SD = 0.66). The next strongest motivational function were Citizen Science (capturing motivations such as contributing to research, engaging with scientists, and better understanding scientific processes, M = 4.03, SD = 0.72), Sociopolitical Responsibility (M = 3.89, SD = 0.95) and Social Motives (M = 3.57, SD = 0.99). The motivation to learn more about the object studied (as assessed with one extra item) was moderately high (M=3.76, SD=1.11). The distribution of scores across these subscales is illustrated in the boxplots (Figure 4B). Correlations between the motivational factors (displayed as indices of the respective items per variable) are provided in the Supplementary Materials (Table S2). Overall motivation (index of all five subscales) was high (M=3.96, SD=.58). A one-way ANOVA revealed significant differences in scores across the six subscales (*F*_5,348_ = 6.41, *p* < 0.001) and post-hoc comparisons showed significant differences among all subscales (Table S3). The MORFEN-CS scale reached high reliability overall as measured with Cronbach’s alpha statistic (α=0.87; α=0.88 including the extra formulated question). Reliabilities per subscale were as follows (Figure 4C): Nature Conservation Values α= 0.75; Social Motives α=0.75; Sociopolitical Responsibility α=0.80; Citizen Science α=0.71; Communication and Organization α=0.44). The effect size, as measured by eta squared (η² = 0.084), indicated a medium-to-large effect. This suggests that participants meaningfully differentiated between distinct types of motivation and that these differences reflect genuine psychological distinctions in what motivated their participation in citizen science.

The survey results highlight several particularly noteworthy findings across subscales (see Table S4). In the domain of Nature Conservation Values, 93.2% of participants reported that they like to advocate for wild organisms (20.3% selecting 4 and 72.9% selecting 5), and 96% indicated with scores 4 or 5 that they aim to take action against the loss of biodiversity. Regarding Sociopolitical Responsibility, nearly half of the respondents (46%) rated engaging in a socially meaningful activity as the highest priority (5). Similarly, 64.4% endorsed (4 or 5) the motivation to initiate societal change through their participation. Findings for the Citizen Science subscale indicate that 78% of participants strongly endorsed (5) the statement that they participate in the project to contribute to a scientific research endeavor. More than half of the participants (54%) endorsed (with 4 or 5) the motivation to better understand scientific processes and to pursue their curiosity about science. Additionally, nearly half of the sample (48%) endorsed (5) that their participation was motivated by the supervision of the project by the Natural History Museum Vienna. Results for an additional survey item (motivation to learn more about *Drosophila*) indicate that the participants engage in the project to learn more about the distribution and ecology of *Drosophila*. For one third of respondents, this motivation was of very high priority (5), around 27% rated it at an intermediate level (4), approximately one third considered it of moderate interest (3), and only 10% indicated lower importance (1 or 2).

### Dissemination

The project achieved broad media outreach through multiple channels, including television, radio, social media, and print. To support the recruitment and engagement of citizen scientists, we produced an introductory video that was published on the NHMW webpage, on YouTube (1,540 followers – 163 views and 9 likes), and on Instagram (21,800 followers – 4,200 views). Over the course of the project, we created additional social media content, published mainly on Instagram and LinkedIn, to promote the project, share progress updates, and encourage further registrations.

After the project was finalized, the Austrian Broadcasting Corporation (ORF) produced a six-minute documentary on its findings for *Mayrs Magazin*, a popular-science television program reaching approximately 3.2 million daily viewers across German-speaking countries (ORF 2021). In addition, science.orf.at, the online platform for science journalism of the ORF, published an article accompanied by a two-minute interview for the ORF radio channel Ö1, which engages approximately 769,000 daily listeners (ORF 2025). The interview was aired twice, and both the documentary clip and the radio interview were made available via the online streaming service of the ORF, further enhancing accessibility and reach.

Beyond broadcast, the findings of the project were also covered in Austria’s leading daily newspapers, reaching an estimated 505,000 readers per edition via *Der Standard*, 421,000 via *Kurier*, and 271,000 via *Die Presse*, respectively. The free newspaper *Heute*, with an extrapolated readership of approximately 675,000 due to its wide distribution (Statistik Austria 2023), served as another key channel to engage participants. Moreover, an article appeared in the weekly Viennese newspaper *Falter*, which prints 46,000 copies per edition (FALTER Medienwelt 2025). These outlets are among the highest-reaching print and hybrid print/online media in Austria, ensuring additional visibility across broad and diverse readerships.

Taken together, this coordinated media presence maximized impact across multiple audiences and communication formats, extending outreach well beyond the immediate project community.

## DISCUSSION

### *Drosophila* research as a suitable model for citizen science projects

In this study we present the citizen science project *Vienna City Fly*, conducted at the NHMW, highlighting both participant engagement and its contribution to urban ecological research using the model system of the genus *Drosophila*. Our findings underscore the dual potential of citizen science emphasized in the introduction: generating valuable ecological data while fostering knowledge, awareness and engagement with urban biodiversity. The project successfully implemented a simple and efficient rotation system to pick-up and deliver fly traps at the NHMW, which was highly accepted and enabled volunteers to collect a total of 18,173 flies across 279 traps, belonging to 13 different *Drosophila* species (Kapun et al. 2025). This demonstrates that citizen scientists can efficiently contribute to comprehensive valuable ecological data collection.

Survey responses (N=59) revealed unique insights into the sociodemographic composition, motivational drivers, levels of engagement, and self-reported impact of participation. The sample was characterised by a relatively high educational background, with three quarters holding at least a master’s degree, and a predominance of female participants, while age distribution was balanced (Figure 2A-C). Recruitment pathways were strongly shaped by social environments of participants, while traditional media and institutional outreach played a smaller role. With 76.3% of participants submitting samples and minimal barriers related to equipment or instructions, the project proved both logistically feasible and scientifically valuable. This aligns with prior research showing that citizen science can expand ecological recording capacities at scales difficult to achieve through professional science alone (Ishida 2020, Olsen et al. 2020). Using *Drosophila* as a research model proved particularly effective: its accessibility and familiarity allowed engagement with minimal prior knowledge while generating data for a largely underexplored field of urban biodiversity research (Kapun et al. 2025).

The project not only succeeded in collecting data, but also generated positive participant outcomes, confirming earlier findings that citizen science can enhance learning and strengthen environmental awareness (Greving et al. 2022, Moczek et al. 2021, Sauermann et al. 2020, Toomey and Domrose 2013). High satisfaction levels and the expressed willingness to participate again highlight its potential as an engaging form of public involvement. At the same time, the request of participants for more scientific content points to opportunities for deepening the educational and dialogic aspects of future initiatives. The reported knowledge gains, the increased positive attitudes towards the study objects, and raised biodiversity awareness further indicate that such projects can extend beyond data collection to foster lasting learning experiences and attitudinal change.

Motivational analyses clarified drivers of engagement. Although motivation appeared high on the subscale Communication and Organization, its very low reliability precludes meaningful interpretation. Accordingly, Nature Conservation Values emerged as the strongest motivator, followed by motivations to contribute to science, Sociopolitical Responsibility, and interest in learning about *Drosophila*. Nearly all participants (97%) reported engaging to contribute to biodiversity conservation, and to participate in socially meaningful work (80%), highlighting the importance of altruistic motives in engaging in citizen science, consistent with prior studies (Kühn et al. 2022, Moczek et al. 2021b).

Institutional trust also played a critical role. High regard for the NHMW underscores the importance of established institutions in legitimizing citizen science and facilitating participation. As discussed above, NHMs uniquely bridge science and society, fostering public engagement, enhancing awareness of environmental change, and supporting conservation efforts (Ballard et al. 2017, Sforzi et al. 2018).

### Potential limitations of the citizen science project

In spite of the overall positive perception of the project by the participants and the demonstrated value of our motivation assessment, several potential limitations of this study need to be acknowledged. First, the findings discussed above must be interpreted with attention to sample composition. Participation in the online survey was voluntarily and therefore likely biased towards participants with positive experiences from the project. Moreover, the sample size and composition was small and not representative of the Austrian population since participants were predominantly highly educated. This suggests that the observed engagement and knowledge gains primarily reflect segments of the population already positively predisposed toward science. While this profile is consistent with findings from earlier citizen science initiatives, it highlights a broader challenge for the field: socially diverse participation remains difficult to achieve (Hobbs and White 2012, Paleco et al. 2021, Pateman and West 2021). This limitation is particularly relevant in the context of democratizing science, since individuals from more marginalized groups – who might benefit most from such initiatives – are often underrepresented. As previous research has shown, and as findings of this study also suggests, recruitment and retention in citizen science tend to favor already science-affine groups (Hobbs and White 2012, West and Pateman 2016), which may limit the inclusiveness and transformative potential of such projects. Consequently, while the project demonstrates that citizen science can reconnect participants with research and biodiversity, its broader societal reach and inclusivity remains limited. Nevertheless, the desire of participants for more scientific content indicates that citizen science can also act as a gateway to deeper engagement with scientific research, potentially multiplying knowledge and awareness within social networks.

Second, while the use of the MORFEN-CS scale represents a strength in grounding this study in an established psychometric framework, the adaptation of the instruments introduces certain methodological limitations. Specifically, the subscale Communication and Organization, that was adapted in this study from its original version, showed insufficient reliability and could not be used for further interpretation. This limitation suggests that the adapted version did not function as robustly as intended and underlines the importance of applying the scale in its original form.

Third, the study relied on self-reported measures of outcomes such as knowledge gain, awareness, attitudes. While these provide valuable insights into perception of participating citizen scientists, they are potentially limited by social desirability bias. Barriers to engagement that were mentioned by participants have also been observed in other citizen science projects (Asingizwe et al. 2020). Addressing such barriers systematically in future work is essential for ensuring data quality and sustaining engagement. Taken together, these limitations highlight the importance of broadening participation, applying validated instruments, and complementing self-reported measures with additional methods that capture behavioral and long-term outcomes when investigating impact of citizen science.

### Future research

Building on the findings of this study, several avenues for future research emerge. First, the motivation to engage in citizen science needs closer examination. A deeper understanding of motivational drivers and related psychological variables – such emotions experienced towards biodiversity and biodiversity loss, pro-environmental behavior and attitudes – is critical for attracting and retaining more diverse audiences. Such knowledge would also enable the strategic design of citizen science projects that maximise both participation and impact.

Second, future work would benefit from the systematic use of validated instruments from social sciences and environmental psychology. Employing established scales allows for comparability between studies and projects, which is currently limited due to heterogeneity in measurement approaches. Standardization in this area would provide a stronger evidence base for assessing the psychological and social effects of participation.

Third, the role of citizen scientists in addressing biodiversity loss and contributing to the achievement of the Sustainable Development Goals (SDGs) deserves further exploration. While the ecological contributions of citizen science are increasingly acknowledged, more empirical research is needed to understand how engagement of participating citizen scientists links to broader sustainability transformations. More generally, incorporating perspectives from behavioral and social sciences is essential for advancing the field. Such approaches can enrich the understanding of how citizen science functions as a form of public engagement with environmental issues and scientific research. They are also central to designing robust impact studies that move beyond self-reported outcomes and capture behavioral change, societal engagement, and long-term effects.

## CONCLUSION

Overall, The Vienna City Fly Project demonstrates that citizen science can generate robust data, raise awareness of urban biodiversity, and provide meaningful motivational experiences. These outcomes meet the growing need for biodiversity data and public engagement (IPBES 2019; Greving et al. 2022). Volunteers collected over 18,000 *Drosophila* specimens from 13 species, contributing valuable ecological insights (Kapun et al. 2025). Nature conservation values and altruistic motives were the strongest drivers of participation, while volunteers reported knowledge gains, more positive attitudes toward *Drosophila*, and increased biodiversity awareness.

In Austria, where public interest in science is declining (Austrian Academy of Sciences 2024), the high satisfaction rates with the project and broad willingness to participate again highlight the potential of low-threshold initiatives to rebuild trust and interest in science. However, the predominance of highly educated participants underscores current limits in inclusivity and societal reach. Natural history museums, as trusted institutions, can play a key role in broadening participation.

Overall, our findings show that *Drosophila* research offers an accessible and effective model system for citizen science, combining ease of participation with strong potential to advance ecological research and public engagement in urban biodiversity.

## DATA AVAILABILITY

The full documentation of the data analysis including the raw data can be found at: https://github.com/capoony/DrosophilaCitizenScience and the raw data table is also available as Supplementary Table S1.

## AUTHORS CONTRIBUTION

### Following the Contributor Role Taxonomy (CRediT; https://credit.niso.org/)

IG: Conceptualisation, Data Curation, Formal analysis, Investigation, Methodology, Validation, Vizualisation, Writing – Original Draft; SS: Conceptualisation, Data Curation, Investigation, Methodology, Validation, Writing – Original Draft; FS: Data Curation, Investigation, Resources, Vizualisation, Writing – Review & Editing; MKi: Project administration, Resources, Writing – Review & Editing; HR: Funding acquisition, Project administration, Writing – Review & Editing; EH: Conceptualisation, Investigation, Validation, Writing – Review & Editing; MKa: Conceptualisation, Data Curation, Formal analysis, Funding acquisition, Investigation, Methodology, Project administration, Software, Supervision, Vizualisation, Writing – Original Draft

## ACKNOWLEDGEMENTS

We are sincerely grateful to the 169 registered and – in particular – the 89 contributing citizen scientists whose dedicated contributions provided the sampling material on which this study is based. We also thank the Horizon Europe-funded FAIRiCUBE project and consortium (grant agreement No 101059238) for financially supporting this project, and in particular the coordinators, Stefan Jetschny and Katharina Schleidt, for their dedicated guidance. Our gratitude extends particularly to our colleague Jana Katharina Köhler at the University of Vienna for her assistance in finding an appropriate psychometric instrument used in this study and for the helpful feedback she provided for this manuscript. We further want to thank the porters at the NHMW for their invaluable help in handling traps – receiving filled ones from citizen scientists and distributing new empty ones. Further, we would like to express our gratitude to Ines Méhu-Blantar for her valuable expertise and support during the initial phase of the citizen science project. Without the generous contributions of all these individuals, and many more at the NHMW, this project would not have been possible.

## Supporting Information

https://github.com/capoony/DrosophilaCitizenScience/tree/main/data

